# DNA microarray analysis of *Staphylococcus aureus* from Nigeria and South Africa

**DOI:** 10.1101/2020.07.22.215632

**Authors:** Adebayo O. Shittu, Tomiwa Adesoji, Edet E. Udo

**Affiliations:** Department of Microbiology, Obafemi Awolowo University, Ile-Ife, 22005; Institute of Medical Microbiology, University Hospital Münster, Domagkstraße 10, 48149 Münster, Germany; Department of Microbiology, Faculty of Medicine, Kuwait University, Safat

**Keywords:** *Staphylococcus aureus*, DNA microarray, antibiotic resistance, virulence, South Africa, Nigeria

## Abstract

*Staphylococcus aureus* is an important human pathogen with an arsenal of virulence factors and a propensity to acquire antibiotic resistance genes. The understanding of the global epidemiology of *S. aureus* through the use of various typing methods is important in the detection and tracking of novel and epidemic clones in countries and regions. However, detailed information on antibiotic resistance and virulence genes of *S. aureus*, and its population structure is still limited in Africa. In this study, *S. aureus* isolates collected in South Africa (n=38) and Nigeria (n=2) from 2001-2004 were characterized using DNA microarray. The combination of *spa* typing and DNA microarray classified the isolates into seven *spa* types and three clonal complexes (CCs) i.e. t064-CC8 (n=17), t037-CC8 (n=8), t1257-CC8 (n=6), t045-CC5 (n=5), t951-CC8 (n=1), t2723-CC88 (n=1), t6238-CC8 (n=1), and untypeable-CC8 (n=1). There was excellent agreement (only two discordant results) between antibiotic susceptibility testing and the detection of the corresponding resistance genes by DNA microarray. Antibiotic and virulence gene markers were associated with specific clones. The detection of the collagen-binding adhesion (*cna*) gene was unique for t037-CC8-MRSA, the enterotoxin gene cluster (*egc*) and staphylococcal complement inhibitor (*scn*) gene for t045-CC5-MRSA, and the combination of genes encoding enterotoxin (*entA*, *entB*, *entK*, *entQ*) were noted with most of the CC8 isolates. The t045-CC5-MRSA clone was positive for the mercury resistance (*mer*) operon. DNA microarray assay provides information on antibiotic resistance and virulence gene determinants and can be a useful tool to identify gene markers of specific *S. aureus* clones in Africa.

## Introduction

*Staphylococcus aureus* is a major human pathogen with a wide array of virulence factors, toxins, and a remarkable ability to acquire antibiotic resistance genes [1,2]. This capability is further enhanced by the constant emergence of new and diverse clones within regions and countries [3]. The knowledge of the epidemiology of *S. aureus*, particularly of methicillin-resistant *S. aureus* (MRSA), is hinged on the application of various typing methods to assist in tracking newly emerging and epidemic clones [4]. Molecular epidemiological typing tools provide valuable information on the emergence of high-risk pandemic *S. aureus* clones, and the prevalence of antibiotic resistance mechanisms and virulence determinants. This is important in the development of intervention strategies and infection control measures in clinical and non-clinical settings [4].

The *S. aureus* epidemiological landscape in Africa has been described mainly through two molecular typing schemes i.e. *Staphylococcus* protein A (*spa*) typing and multilocus sequence typing (MLST) [5,6]. These studies revealed that the most widely distributed MSSA clones in Africa include ST5, ST8, ST15, ST30, ST121, and ST152. Whereas ST5, ST30, ST121, and ST152 are predominant in Central and West Africa, ST8, ST15, ST30 are dominant in North Africa [5]. As for MRSA, ST239/241 is dominant in many African countries, ST8 and ST88 in West, Central and East Africa, ST80 in North Africa, and ST5, ST36 and ST612 in South Africa [5,6]. However, data on the repertoire of antibiotic resistance and virulence genes of *S. aureus*, and its clonal diversity in Africa are limited. In this study, we characterized archived *S. aureus* isolates from Nigeria and South Africa using DNA microarray. The study aimed to provide detailed information on antibiotic resistance and virulence-related genes, and the population structure of the isolates. This could provide information on antibiotic resistance and virulence genes that could represent epidemiological markers to specific *S. aureus* clones in Africa.

## Materials and methods

### Bacterial isolates

The *S. aureus* isolates have been described in previous investigations [7,8] and were obtained from different clinical samples from 2001-2004. They comprised mainly *S. aureus* from South Africa (MRSA: n=37; MSSA: n=1), while two isolates from Nigeria were included based on their resistance to cefoxitin and mupirocin.

### *Spa* typing and DNA microarray

*Spa* typing was performed by sequencing of the hyper-variable region of the protein A gene (*spa*), as described previously [9]. Further characterization of the isolates was performed using the DNA microarray technique [10]. This assay detects various genetic determinants including species markers, genes encoding resistance and virulence, toxin and immune evasion complex, and the arginine catabolic mobile element (ACME). Others include adhesion and biofilm genes, microbial surface components recognizing adhesive matrix molecules (MSCRAMMs), accessory gene regulator (*agr*), capsule and SCC*mec* types. Furthermore, this assay can also delineate *S. aureus* to sequence types (STs) and/or clonal complexes (CCs). The analysis was performed using the Identibac *S. aureus* genotyping Kit 2.0 (Alere Technology, Jena, Germany) as described previously [10]. The isolates were classified based on the *spa* type and clonal complexes (*spa*-CC).

## Results

The combination of specific *S. aureus* markers confirmed the identity of the isolates (n=40) (Supplementary material). Based on the microarray data, all the isolates harboured genes encoding proteases (*splA*, *splB*, *sspA*, *sspB* and *sspP*), MSCRAMMs (*bbp*, *clfA*, *clfB*, *ebpS*, *fib*, *fnbA*, *map*, *sasG*, *sdrC*, *vwB*), leukocidin (*lukF* and *lukE*), haemolysin (*hlgA*), and intracellular adhesion (*icaA*). However, none possessed the exfoliative toxin (*etA*, *etB*, *etD*), epidermal cell differentiation (*edinA, edinB* and *edinC*), surface protein involved in biofilm production (*bap*), and the ACME genes (Table 1). Only two discordant results were observed between the antibiotic susceptibility testing and the detection of the corresponding resistance genes (Table 2). A penicillin-resistant isolate was *blaZ* gene negative, while an isolate susceptible to erythromycin possessed the *ermC* gene.

Molecular typing classified the isolates into seven *spa* types t037, t045, t064, t951, t1257, t2723 and t6238, and three clonal complexes (CCs), CC5, CC8 and CC88. The delineation of the various groups (*spa*-CC) and their unique characteristics are described (Table 2).

### CC5

#### t045-CC5 (South German EMRSA or the South German EMRSA/Italian Clone)

Five MRSA isolates belonged to t045. They were grouped with *agr* group II and capsule type 5. While most of them (4/5) possessed the SCC*mec* II element, the cassette chromosome recombinase genes A/B-2 was not detected in one MRSA isolate and was assigned to SCC*mec* type I (Supplementary material). All the t045 isolates harboured the resistance genes for aminoglycosides (*aacA-aphD* and *aphA3*), macrolide (*ermA*), fosfomycin (*fosB*), streptothricine (*sat*), and quaternary ammonium compounds (*qacA*). Besides, they were positive for the mercury resistance operon (*mer*). A unique feature of this clone was the detection of the enterotoxin gene cluster (*egc*), the presence of only one of the immune evasion cluster (IEC) genes (*scn*), and the non-detection of the tetracycline resistance genes (*tetK*, *tetM*).

### CC8

The CC8 isolates belonged to five *spa* types, consisting of t064 (n=17), t037 (n=8), t1257 (n=6), t951 (n=1) and t6238 (n=1). They were associated with *agr* group I and capsule type 5, except those belonging to *spa* type t037 that belonged to capsule type 8. One MRSA each could not be characterized by *spa* and *agr* typing.

#### t037-CC8 (Vienna/Hungarian/Brazilian clone)

The t037 *spa* type was represented by eight isolates. They possessed the SCC*mec* type III genetic element as well as the *mer* and the recombinase (*ccrC*) genes (Supplementary material). Furthermore, all the t037 isolates possessed the *aphA3* and *tetM* genes, and those that exhibited phenotypic resistance to erythromycin (n=8) and chloramphenicol (n=4) were positive for the corresponding genes (*ermA* and *cat*) (Table 2). Seven isolates harboured the fosfomycin and streptothricine resistance determinants (*fosB*, *sat*). The *aacA-aphD* and *tetK* genes were identified in at least five isolates that were resistant to gentamicin and tetracycline, respectively. All the trimethoprim-sulphamethoxazole-resistant MRSA were *dfrA* (dihydrofolate reductase) negative. The enterotoxin genes (*entA*, *entK* and *entQ*) were detected in at least six isolates, while the distinctive feature of this clone was the positive result for the collagen-binding adhesion (*cna*) gene.

#### t064-CC8 (USA500)

This clone comprised 16 isolates (MSSA n=1; MRSA n=15) from South Africa and MRSA (n=1) from Nigeria. The SCC*mec* type IV was identified in all the MRSA from South Africa, while the isolate from Nigeria carried the SCC*mec* V element and the *mer* operon (Table 2). The following genes i.e. *aacA-aphD*, *dfrA, ermC*, and *tetM* were detected in at least 14 of the 17 isolates. Only two MRSA were *qacA*-positive. The combination of genes encoding enterotoxin (*entA*, *entB*, *entK*, *entQ*) was a common feature noted with most of the isolates.

#### t951-CC8 (Lyon Clone/UK-EMRSA-2)

The only MRSA (SCC*mec* IV) possessed the antibiotic resistance genes (*tetM* and *fosB*), and enterotoxin A gene. It was also positive for the immune evasion cluster genes (*sak* and *scn*).

#### t1257-CC8 (USA500)

The six isolates belonging to this *spa* type were associated with SCC*mec* type IV. Moreover, the isolates exhibited similar antibiotic resistance gene profiles (*aacA-aphD, dfrA, ermC*, and *tetM*), enterotoxin (*entA*, *entB*, *entK*, *entQ*) and immune evasion (*sak*, *scn*) gene content with those assigned with t064-CC8.

#### t6238-CC8 (USA500)

The single isolate associated with this *spa* type harboured the SCC*mec* IV element, and the antibiotic resistance (*aacA-aphD, dfrA*, *fosB*, *tetM*) and enterotoxin (*entA*, *entB*, *entK*) genes were identified. Furthermore, the isolate was only positive for one of the IEC genes (*sak*). It was negative for *agr* types 1-IV.

#### *spa* untypeable-CC8 (USA500)

The gene content of the MRSA isolate was similar to other members of CC8.

#### t2723-CC88

A single MSSA isolate was associated with t2723. It was assigned to *agr* group III and capsule type 8. Phenotypic resistance to tetracycline and mupirocin was confirmed by the detection of the *tetK* and *mupR* genes, respectively. No enterotoxin gene was detected in this isolate. However, it was positive for the IEC (*chp*, *sak*, *scn*) and Panton-Valentine Leukocidin (PVL) genes.

## Discussion

The combination of *spa* typing and DNA microarray was utilized to characterize *S. aureus* isolates obtained in South Africa and Nigeria. The DNA microarray is a DNA-DNA hybridization method containing several probes for the rapid identification, characterization of *S. aureus* resistance and virulence gene profiles, and their assignment into clonal complexes [10]. The results showed excellent correlation between antibiotic susceptibility testing and the corresponding resistance gene profiles by DNA microarray. The results also revealed the association of some antibiotic resistance gene determinants with certain MRSA clones. Specifically, the *aphA3*, *ermA*, and *mer* genes were unique characteristics associated with t037-CC8-MRSA and t045-CC5-MRSA (Table 2). Similarly, although these two clones exhibited resistance to trimethoprim-sulphamethoxazole, they were negative for *dfrA* that encodes resistance to trimethoprim in *S. aureus*. Although trimethoprim resistance in *S. aureus* can be due to any of three determinants, *dfrA*, *dfrG* and *dfrK*, the *dfrG* is associated with trimethoprim resistance in the majority of the trimethoprim-resistant *S. aureus* in Africa [11,12]. Trimethoprim-resistant *S. aureus* isolates harbouring *dfrG* and associated with *spa* types t037 and t064 have also been reported in Nigeria [11], which is similar to our findings in this study. This observation suggests that *dfrG* was responsible for trimethoprim resistance in our *dfrA*-negative isolates.

Interestingly, although t037-CC8-MRSA and t045-CC5-MRSA shared common antibiotic resistance determinants, they differed in the carriage of the tetracycline resistance determinants (*tetK*, *tetM*) that was present in t037-CC8-MRSA and not in t045-CC5-MRSA (Table 2). MRSA is characterized by the presence of the staphylococcal cassette chromosome *mec* (SCC*mec*), a mobile 21-to 60-kb genetic element, and 13 SCC*mec* types have been identified [13]. The SCC*mec* types II (53.0 kb) and III (66.9 kb) are large elements due to the acquisition and insertion of mobile genetic elements (MBEs). The antibiotic resistance genes observed in the two clones have been identified on MBEs such as transposons including *Tn*554 (*ermA*), *Tn*4001 (*aacA-aphD*), *Tn*5405 (*aphA3*, *sat*), *Tn*916 (*tetM*), and plasmids i.e. pT181 (*tetK*), pI258 (*mer*) and pNE131 (*ermC*) [14,15,16]. The t037-CC8-MRSA and t045-CC5-MRSA lineages are typical hospital-associated clones, and their multi-resistant nature are attributed to the various MBEs that harbour different antibiotic resistance genes in the joining regions J1 to J3 [17].

We note with interest that all the t045-CC5-MRSA isolates possessed the mercury resistance operon, a feature also commonly present with t037-CC8-MRSA. The mechanism for the acquisition of SCC*mercury* by *S. aureus* is still unclear although two views have been postulated. The first suggests that this gene determinant may have been integrated into an SCC element with the emergence of SCC*mercury* in coagulase-negative staphylococci, which is subsequently transferred to *S. aureus*. The second opinion is that a plasmid (e.g. pI258) harbouring the resistance gene determinant to the quaternary ammonium compound could have been transferred to *S. aureus* and integrated into an SCC element to form SCC*mercury* [18]. Interestingly, a comparison of our results with a previous report consisting of a wide collection of CC5-MRSA isolates in the Western Hemisphere [19] revealed that the presence of *mer* gene in t045-CC5-MRSA is a rare feature of this clone. Therefore, future studies should ascertain whether this observation represents a recent acquisition of the *mer* operon by CC5-MRSA. SCC*mec* types IV (20.9-24.3 kb) and V (28 kb) typically encode resistance only to β-lactam antibiotics. We observed that, in addition to β-lactam resistance, the isolates classified as t064/t1257/t6238-CC8 harbouring the SCC*mec* IV element also possessed genes (*aacA-aphD*, *dfrA*, and *ermC*) mediating resistance to aminoglycosides, trimethoprim and macrolides, respectively. The presence of these resistance determinants in our archived isolates support existing data [20,21] that this multi-resistant lineage is established and well adapted in the hospital environment in South Africa.

Mupirocin is a topical antibiotic that is widely used for nasal decolonization and prevention of *S. aureus* infections. However, the emergence and increasing rates of resistance and treatment failure are major drawbacks [22]. Two levels of mupirocin resistance have been elucidated i.e. low-level and high-level resistance attributed to various chromosomal mutations, and the acquisition of plasmids (harbouring *mupA* or *mupB* genes), respectively [23,24]. Decolonization is ineffective with patients and personnel colonized with high-level mupirocin resistant (mupR) MRSA [22]. Moreover, mupirocin resistance could also facilitate the spread of multidrug resistance through co-selection with other plasmid-borne resistance genes [25,26] In this study, the genetic background of a high-level mupR MSSA was determined (PVL-positive, t2723-CC88). Only two studies have provided information on the genetic lineage of high-level mupR *S. aureus* from clinical samples in Africa which include t127, t4805 (MSSA), and t032, t1467 (MRSA) [27,28]. The prevalence and burden of mupR *S. aureus* are still unclear in many countries in Africa [29]. CC88-MRSA is an established lineage in West, Central and East Africa [5], and the identification of a *mupA*-PVL-positive MSSA from this background is worthy of note. Prospective national and continental studies are important to evaluate the prevalence, burden and genetic background of mupR *S. aureus* in Africa.

*S. aureus* produces a range of virulence determinants including at least 23 exotoxins which are categorized into staphylococcal enterotoxins (SEs) comprising SEA-SEE, and staphylococcal enterotoxin-like (SEl) consisting of SEG-SElY [30]. They belong to the family of superantigens (SAgs) with a unique feature to act primarily on the intestine to cause enteritis characterized by emesis [31]. Our investigation indicated that some enterotoxin genes were associated with specific genetic backgrounds, which is in support of previous reports [32,33]. The t037-CC8-MRSA was characterized by the detection of *entA*, *entK*, and *entQ* genes. The egc cluster (*entG*, *entI*, *entM*, *entN*, *entO*, and *entU*) were associated with t045-CC5-MRSA, while the *entA*, *entB*, *entK* and *entQ* genes were linked with t064/t1257-CC8. The SE genes are carried and disseminated through different MBEs which include prophages, plasmids, transposons, and *S. aureus* pathogenicity islands (SaPIs) [30]. The *entA*-*entK*-*entQ* genes is found on the prophage ΦSa3ms and ΦSa3mw, the egc cluster on the genomic island vSaβ, and *entB*-*entK*-*entQ* have been identified on SaPI3 [34].

## Conclusions

This study characterized archived *S. aureus* isolates from Nigeria and South Africa using two molecular-based typing methods (*spa* typing and DNA microarray). A high level of agreement was observed between antibiotic susceptibility testing and the identification of the corresponding gene determinants using DNA microarray. Also, some antibiotic resistance and virulence genes were associated with specific clonal lineages. The *aphA3*, *ermA*, and *mer* genes were associated with hospital-associated clones (t037-CC8-MRSA and t045-CC5-MRSA), *cna* with t037-CC8-MRSA, the *egc* cluster and *scn* with t045-CC5-MRSA, and *entA*, *entB*, *entK*, *entQ* with most of the CC8 isolates. There are some limitations to this study. These include the small number of *S. aureus* isolates analyzed, and the limitations of the DNA microarray technology i.e. high cost of reagents and equipment in resource-limited settings, cross-hybridization reaction, and a moderate level of reproducibility. Nevertheless, the main advantages of the technology include speed compared with procedures involving several PCR and gel electrophoresis, the diverse array of genes investigated, and the quantum of data generated. DNA microarray analysis has provided useful information on gene determinants for antibiotic resistance and virulence, and their relationship with some *S. aureus* genetic background in Nigeria and South Africa. Although the outcome of this investigation is not representative of the diverse *S. aureus* clonal lineages in Africa, the genetic markers noted could be a useful adjunct in the molecular typing and tracking of new and emerging *S. aureus* clones on the continent.

## Acknowledgements

This study received support from the Deutsche Forschungsgemeinschaft (SCHA 1994/5-1) and the Alexander von Humboldt Foundation (“Georg Forster-Forschungsstipendium” which was granted to AOS). We appreciate the technical assistance of Mrs Tina Verghese and Bindu Mathew.

## Authors Contributions

Conceptualization: Adebayo O. Shittu, Edet E. Udo.

Data Curation: Adebayo O. Shittu, Edet E. Udo.

Funding acquisition: Adebayo O. Shittu, Edet E Udo.

Investigation: Adebayo O. Shittu, Edet E. Udo.

Methodology: Adebayo O. Shittu, Edet E. Udo.

Project Administration: Edet E. Udo.

Supervision: Edet E Udo.

Writing – original draft: Tomiwa Adesoji, Adebayo O. Shittu.

Writing – review & editing: Adebayo O. Shittu, Tomiwa Adesoji, Edet E Udo.

## Ethical standards

Ethical clearance was not necessary as only isolates were analyzed in this study.

## Supporting information

Detailed characteristics of the *S. aureus* isolates (n=40) including antibiotyping, *spa* typing and DNA microarray hybridization results of antibiotic and virulence genes.

